# Deterministic processes played more important roles than stochastic processes in driving community assembly of ectomycorrhizal fungal associated with *Pinus tabuliformis –* an endemic Pinaceae species in China

**DOI:** 10.1101/2024.07.05.602164

**Authors:** Yongjun Fan, Zhimin Yu, Jinyan Li, Xuan Zhang, Min Li, Jianjun Ma, Yonglong Wang

**Affiliations:** Yinshanbeilu Grassland Eco-hydrology National Observation and Research Station, China Institute of Water Resources and Hydropower Research; School of Life Science and Technology, Inner Mongolia University of Science and Technology, Baotou, Inner Mongolia, China; College of Ecology and Environment, Baotou Teacher’s College, Baotou, Inner Mongolia, China; College of Life Science and Technology, Inner Mongolia Normal University, Hohhot Inner Mongolia, China; College of Life Science, Lang Fang Normal University, Lang fang, Hebei Province, China

**Author notes:** Corresponding Author: Yonglong Wang, No. 3 Science Road, Qingshan District, Baotou, Inner Mongolia, 014030, China, Email address, Jianjun Ma, No. 100 Aimin West Road, Langfang City, Hebei Province, 065000, China.

**Keywords:** Fungal diversity, *Pinus tabuliformis*, Community assembly, Deterministic process, Inner Mongolia

## Abstract

*Pinus tabuliformis* represents a typical and essential woody ectomycorrhizae (EM) host plant growing in North China. EM fungi contribute to the host healthy and the stability of the forest ecosystem. However, EM fungal community associated with this tree is less documented. We examined EM fungal diversity and composition of *P. tabuliformis* from three sites in Inner Mongolia, China by using Illumina MiSeq sequencing on the rDNA ITS2 region. Our results showed that a total of 105 EM fungal operational taxonomic units (OTUs) were identified from 15 composite root samples, and the dominant lineages were /*suillus-rhizopogon*, /*tomentella-thelephora*, /*tricholoma*, /*amphinema-tylospora*, /*wilcoxina*, /*inocybe* and /*Sebacina*. A high proportion of unique EM fungal OTUs were detected and some abundant OTUs preferred to exist in specific sites. EM fungal communities were significantly different among the sites, with soil, climatic and spatial variables being responsible for the community variations. The EM fungal community assembly was mainly governed by deterministic processes. These findings suggest that this endemic Pinaceae species in China also harbored a rich and distinctive EM fungal community and deterministic processes played more important roles than stochastic in shaping the symbiotic fungal community. Our study improves our understanding of EM fungal diversity and community structure from the perspective of a single host plant that has not been investigated exclusively before.

## INTRODUCTION

As the symbiont formed between soil fungi and plant, ectomycorrhiza (EM) played critical roles in forest ecosystem, such as affecting the host plant healthy, promoting nutrients and water mobilization between individuals of the same or different host plant species via belowground mycelial networks, and thus contributing to plant diversity, ecosystem functions and stability (Smith and Read, 2008; Tedersoo et al., 2020; Anthony et al., 2022; Averill et al., 2022; Cahanovitc et al., 2022). With the rapid development of sequencing techniques, particularly high-throughput sequencing, EM fungal diversity and community at diverse forest ecosystems and various spatial scales received much attention (Tedersoo et al., 2013a; Wang et al., 2019; Gong et al., 2022; Miyamoto et al., 2022). Researchers explored the EM fungal diversity and variables responsible for community structure. Commonly, the EM fungal community was affected by host plant identity or phylogeny (Wu et al., 2018; Wang et al., 2021a), spatial scales (Glassman et al., 2015; Wang et al., 2021b), habitat types (Mundra et al., 2016; Gong et al., 2022; Miyamoto et al., 2022; Yang et al., 2022), soil and climatic factors (Miyamoto et al., 2015, 2018; Glassman et al., 2017; Kwatcho et al., 2022), and also host age (Gao et al., 2015; Mandolini et al., 2022). For example, host plant phylogeny explained the largest variations in EM fungal community associated with Betulaceae, Fagaceae, and Salicaceae plants, and in the permafrost ecosystem of eastern Siberia (Tedersoo et al., 2013a; Wu et al., 2018; Wang et al., 2019; Miyamoto et al., 2022), while climatic conditions dominantly shaped EM fungal communities on two Japanese mountains apart from ∼550 km (Miyamoto et al., 2015); Certainly, spatial distance also participate in predicting EM fungal community at scales ranging from local to the global (Glassman et al., 2015; Wang et al., 2019a; Prieto-Rubio et al., 2022). Thus, many previous studies investigated variables responsible for EM fungal community. Still, it is evident that the main drivers in determining EM fungal community varied in studies conducted at various scales and backgrounds.

Revealing the mechanisms underlying microbial community assembly is critical to gain a clearer understanding of biodiversity maintenance and predicting the response of microbes to global changes. Commonly, microbes suffer from the selection pressure derived from biotic and abiotic factors (i.e., environmental filtering), because different fungal species preferred specific environmental conditions such as plant, soil, and climatic, which can be regarded as deterministic processes (Vellend, 2010). Meanwhile, dispersal limitation, drift, and speciation also affected the microbial community, which can be defined as stochastic processes (Stegen et al., 2012; Zhou et al., 2017; Chen et al., 2019). It has been widely accepted that both deterministic and stochastic processes are important in driving microbial community assembly, but the relative importance of the different processes on community assembly of microbes varied with spatial scales, habitats, and microbial groups (Zheng et al., 2021; Prieto-Rubio et al., 2022; Wang et al., 2022). For instance, soil fungal communities in 18 oceanic islands of China were mainly predicted by deterministic processes (Zheng et al., 2021); Stochastic processes dominantly drove soil fungal communities along an altitudinal gradient on Tibetan plateau (Hussain et al., 2021); community assembly of bacteria in natural mountain forests of eastern China was mainly predicted by deterministic processes (Ni et al., 2021). However, the mechanisms underlying the community assembly of EM fungi remain unclear now.

*Pinus tabuliformis* is an endemic pine species in North China, with high ecological and economic values as it can provide timbers and be used for afforestation, and it is also the dominant EM woody host plant in temperate forests. However, we know less about EM fungal diversity, community composition, and mechanisms underlying the community assembly of this EM host plant. Thus, under this environmental context, we tried to describe the EM fungal community structure of *P. tabuliformis* in Inner Mongolia, China. The Illumine high-throughput sequencing was adopted to sequence the EM fungi of root samples collected from three sites in Inner Mongolia. We sought to (1) explore EM fungal diversity and community composition of *P. tabuliformis* and (2) reveal the mechanisms underlying the EM fungal community assembly of *P. tabuliformis*. We hypothesized that *P. tabuliformis* harbor high and some distinct EM fungal groups; the deterministic processes played more critical roles in driving the community assembly of EM fungi associated with *P. tabuliformis*.

## MATERIALS AND METHODS

### Study sites and sample collection

This study was conducted in three typical forests in Inner Mongolia, China (**Fig. S1**). The mean annual temperature (MAT) ranged from 2.42-5.97 °C, and mean annual precipitation (MAP) ranged from 312-483 mm, according to the data extracted from the WorldClim dataset at the resolution of 30-arcsecond (Hijmans et al., 2005). In the forests, *P. tabuliformis* was the dominant woody EM host plants, with some other EM hosts such as Betulaceae and Salicaceae plants. The sampling work was conducted on August 2018, which is the summer season. At each site, five individuals were selected for sampling. The individuals are at least 10 m apart to ensure sample independence. The fine roots were carefully excavated at three points of each individual and merged as one composite sample. The root samples were transported to the laboratory within 24 hours in an ice box and stored at t −80 ◦C until molecular analysis. Meanwhile, five soil samples were collected at each site when root sampling, and were merged as one composite sample. The soil samples were air-dried, sieved by 2 mm mesh, and then used to analyze soil properties. Thus, a total of 15 root samples and three soil samples were obtained from the three sites in our study. The geographic coordinates and altitude were recorded by using a high-sensitivity GPS instrument. Details on geographic and climatic information of each site are available in **Table S1**.

### Soil properties analysis

Soil pH was determined using dried soil mixed with distilled water at a 1:2.5 ratio (w/v) using a digital pH meter (Mettler Toledo, Zurich, Switzerland). The total carbon (C) and total nitrogen (N) were measured by direct combustion using a Vario EL III C/N Element Analyzer (Elementar Analysensysteme GmbH, Germany). Total phosphorus (P), total potassium (K), and total calcium (Ca) were assessed using an iCAP 6300 inductively coupled plasma spectrometer (Thermo Scientific, Wilmington, USA). A detailed descriptions of the soil properties analysis can be found in Wang et al. (2019).

### Molecular analysis

The roots were firstly washed using tap water and cut into 1-2 cm fragments. The EM root tips were identified according to specific morphological characteristics (e.g. color and emanating hyphae) under a stereomicroscope. About 200 healthy EM root tips were randomly picked from each sample, thus resulting in 30,000 root tips used for analysis in our study. The EM root tips were cleaned carefully with sterilized distilled water and stored at −80 ◦C until total DNA extraction.

The total DNA extraction was conducted using the CTAB method. A semi-nested PCR was adopted to amplify the fungal ITS2 region. Primers of ITS1F and ITS4 were first used in the first round of PCR to amplify the entire ITS region, and the primers of fITS7 and ITS4 were used to target the ITS2 region. The primer ITS4 used in the second PCR amplification was equipped with unique barcode sequence to distinguish each sample in the bioinformatic analysis. Detailed information on PCR step was available in Wang et al. (2021). Three PCR replicates were conducted for each sample and then pooled to generate a composite PCR product. After the amplification, the PCR products of each sample were purified using the Wizard SV Gel and PCR Clean-Up System (Promega, Madison, USA). The amplicon concentration of each sample was determined using nanodrop spectrophotometer (Nanodrop 2000, NanoDrop Technologies, Wilmington, USA). Thereafter, the purified amplicons were pooled with an equimolar amount (100 ng) for each sample. The sequencing work was performed on an Illumina MiSeq PE300 platform using the paired-end (2 × 250 bp) option at the Environmental Genome Platform of Chengdu Institute of Biology, Chinese Academy of Sciences, China.

### Bioinformatic analysis

Clean data was generated from raw data after quality control using the QIIME platform (Caporaso et al., 2010). The ITS2 region was extracted using the ITSx software (Bengtsson-Palme et al., 2013). The chimera sequences were detected and removed by referring the UNITE database (Kõljalg et al., 2013). After that, the high-quality ITS2 sequences were clustered into operational taxonomic units (OTU) based on 97% similarity using the UPARSE pipeline after de-singleton and de-replicates (Edgar 2013). The representative sequence (most abundant) of each OTU was searched against UNITE database (Kõljalg et al., 2013) using the basic local alignment search tool (BLAST). The taxonomy of each OTU was analyzed based on criteria proposed by Tedersoo et al. (2014). After this, the EM fungal OTUs were identified according to Tedersoo and Smith (2013) if they matched the known EM fungi and lineages. To eliminate the effect of heterogenous sequence depth among samples on following community analysis, we rarefied the EM fungal reads to 1,542 per sample by using the rrarefy command in the vegan package (Oksanen et al., 2007). Raw sequences have been deposited in the sequence read archive of NCBI under accession no. PRJNA902691.

### Statistical analysis

All data analyses were conducted in R v. 4.0.1 (R development core team, 2021). Species accumulation curves of observed EM fungal OTUs in each site were drawn using the specaccum command in the vegan package (Oksanen et al., 2007). Diversity index including OTUs richness, Shannon index, and Simpson index were calculated using the diversity command in the vegan package (Oksanen et al., 2007), and then tested for normality and the homogeneity of variance before analysis of variance (ANOVA). One-way ANOVA was performed to examine the effect of the site on the fungal diversity index mentioned above. The interactive Venn diagram and Venn network were generated using the online software Evenn (Chen et al., 2021) to analyze the unique and shared EM fungal OTUs across the three sites. Based on the latitude and longitude, the pairwise Euclidean distance among the three sites was calculated, which were then converted into the principal coordinates of neighbor matrices (PCNM) eigenvector using the pcnm command in the PCNM package (Dray et al., 2006). Generalized linear models (GLMs) with Poisson distribution were constructed to test the effect of spatial, climatic, and soil variables on EM fungal OTUs richness. The multi-model inference approach was applied to obtain the best GLM model, which was generated from a set of reduced models by using dredge and model.avg commands in the MuMIn package (Barton 2018).

A Krona chart showing the different taxonomic levels of the EM fungal community was generated using the online Krona tool (v.2.6, Ondov, Bergman & Phillippy, 2011). A distance matrix for EM fungal community (Hellinger-transformed data) was constructed by calculating dissimilarities using the Bray-Curtis method. Non-metric multi-dimensional scaling (NMDS) analysis was conducted to visualize the difference in the EM fungal communities among the three sites using the metaMDS command in the vegan package (Oksanen et al., 2007). To identify the significant factors that affect the EM fungal community, spatial PCNM eigenvector, climatic and soil variables were fitted into the NMDS plot using the envfit command in the vegan package (Oksanen et al., 2007). Permutational multivariate analysis of variance (PerMANOVA) was adopted to determine the significance of the difference in the fungal communities among sites using the adonis command in the vegan package based on 999 permutations (Oksanen et al., 2007). Additionally, random forest analysis was performed to identify the significant variables responsible for the fungal community composition by using the randomForest command in the randomForest package (Liaw and Wiener, 2019). The importance and significance of each variable were assessed by using the rfPermute and rp.importance command in the rfPermute package (Archer, 2013), and the rf.significance command in the rfUtilities package was used to test for model significance (Evans & Murphy, 2018).

Relative abundances of the abundant fungal OTUs (> 0.5% of total reads) among the sited were displayed by using the pheatmap command in the pheatmap package (Kolde 2015). Additionally, in order to investigate the distribution difference of EM fungal OTUs across sites, preference analysis designed by Toju et al. (2016) was performed based on the site-OTUs matrix, in which rows represented sampling sites, columns represented fungal OTUs, and the cell entries indicated the numbers of samples of specific site-fungus combinations. The detailed description of this analysis method was available in Toju et al. (2016). To quantify the relative importance of the stochastic and deterministic processes to community assembly, the modified normalized stochasticity ratio (MST) index was calculated using the tNST command in the NST package (Ning et al., 2019). The MST index ranges from 0 to 100%, with 50% as the boundary point between more stochastic (>50%) and more deterministic (<50%) assemblies. To increase the robustness of the MST analysis, two distance metrics (Bray and Jaccard) coupled with two null model algorithms of Taxa-Richness constraints of proportional-proportional (PP) and proportional-fixed (PF) were adopted in the analyses.

## RESULTS

### Fungal database summary

A total of 288,458 high-quality ITS2 sequences were generated from 290,326 raw sequences after quality filtering. The high-quality ITS2 sequences were assigned into 299 OTUs, of which 136 OTUs were identified as EM fungi. After rarefying all samples to the same sequencing depth (1,542), 105 EM fungal OTUs were retained for the subsequent statistical analyses. The 28 most abundant OTUs (>0.5%) occupied 94.4% of the total EM fungal sequences (**Fig. S2A**), and 81 out of 105 (77.1%) EM fungal OTUs occurred in less than three samples (**Fig. S2B**).

### EM fungal diversity

The accumulation curves of the observed EM fungal OTUs in each site did not reach the plateau, indicating that further sampling could result in more undiscovered EM fungi (**Fig. 1A**). The EM fungal OTUs richness ranged from 11.2±2.1 (Mean±SE) in WLS to 16.2±8.8 in CF and 16.4±2.5 in HLH. One-way ANOVA indicated that the diversity indices (observed EM fungal OTUs richness, Shannon and Simpson values) were not significantly different among the sites (**Fig. 1B-D**). Venn diagrams showed that CF harbored the highest proportion of unique OTUs (33, 31.4% of total OTUs), followed by HLH (28, 26.7%) and WLS (14, 13.3%); Meanwhile, only eight OTUs (7.6%) were shared by the three sites (**Fig. 2**). GLM result indicated that MAP, MAT, soil K, and altitude are the best predictors of EM fungal OTUs richness (**Table 1**).

**Fig. 1.**
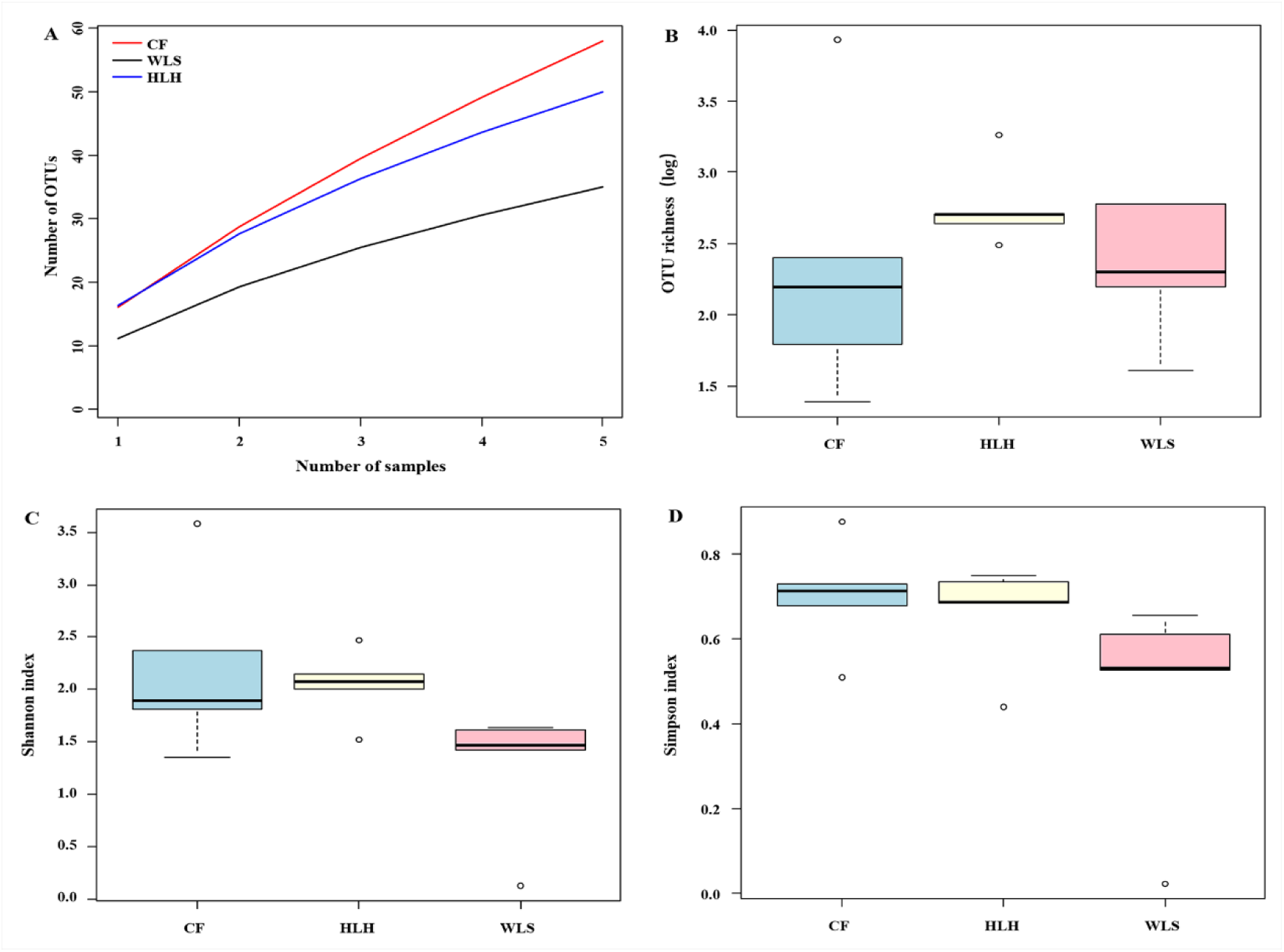
Ectomycorrhizal (EM) fungal diversity. Accumulation curves of EM fungal operational taxonomic units (OTUs) (A), EM fungal OTUs richness (B), Shannon index (C) and Simpson index (D) at three sites. CF, Chifeng; HLH, Heilihe; WLS, Wulashan.

**Fig. 2.**
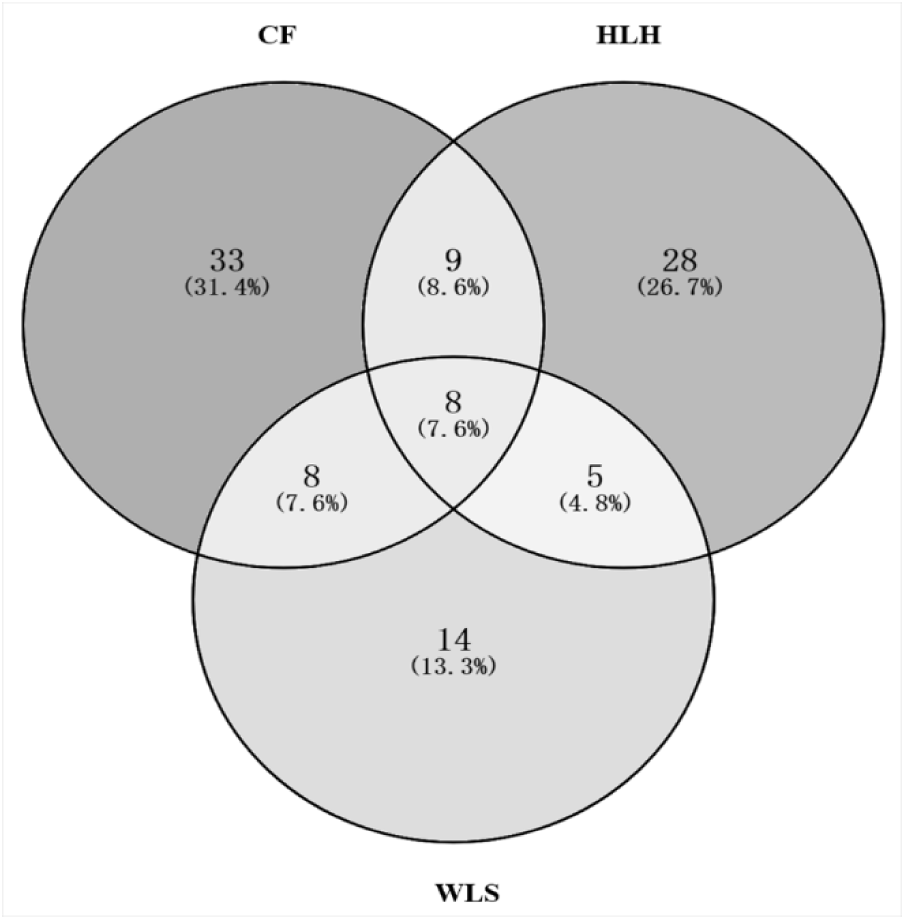
Venn diagram showing the shared and unique ectomycorrhizal fungal operational taxonomic units (OTUs) of the three sites. Deeper color represents higher proportion of OTUs. CF, Chifeng; HLH, Heilihe; WLS, Wulashan.

**Table 1.** Generalized linear model revealed significant variables responsible for ectomycorrhizal fungal operational taxonomic unit richness.

|  | Estimate | SE | z value | P value |  |
| --- | --- | --- | --- | --- | --- |
| (Intercept) | 2.47 | 0.60 | 4.05 | <0.001 | *** |
| MAP | <0.01 | <0.01 | 2.20 | 0.03 | * |
| K | -0.03 | 0.01 | 2.19 | 0.03 | * |
| MAT | 0.11 | 0.05 | 2.17 | 0.03 | * |
| Altitude | <0.01 | <0.01 | 2.16 | 0.03 | * |

MAP, mean annual precipitation; K, soil total potassium; MAT, mean annual temperature.

The Krona diagram showed that *Tomentella*, *Tricholoma*, *Rhizopogon*, *Amphinema*, *Suillus*, *Wilcoxina*, *Inocybe*, and *Geopora* were the dominant fungal genera, accounting for 90% of total fungal sequences (**Fig. 3**). All EM fungal OTUs were assigned into 21 lineages, of which /suillus-rhizopogon, /tomentella-thelephora, /tricholoma, /amphinema-tylospora, /wilcoxina, /inocybe and /Sebacina, accounting for 90.4% of total sequences (**Table 2 and Fig. 4**). NMDS ordination showed that the EM fungal communities were clearly separated between each other (**Fig. 5**), and the PerMANOVA further demonstrated that EM fungal communities were significantly different among sites (adonis: *R*^2^ = 0.30, *P* = 0.001). The environmental fitting test indicated that the spatial, climatic, and soil variables were significantly correlated with EM fungal community structure (**Fig. 5 and Table S2**). Similarly, random forest analysis indicated that all spatial, climatic, and soil variables were significant predictors of EM fungal community structure, and explained 18.87% of the variations in communities (**Fig. 6**). At the OTU level, the heatmap showed that the relative abundance of some abundant OTUs was different among the three sites (**Fig. S3**). Site/fungus preference analysis indicated that all sites harbored the EM fungal OTUs that showed site preference, 11 out of 41 (26.8%) abundant EM fungal OTUs showed significant preference to specific site, and 14 of 123 pairs of site-fungal OTUs exhibited remarkably strong preferences (**Fig. 7**). The MST model showed that the MST values of all sites were below 50%, indicating the EM fungal communities were mainly shaped by the deterministic processes (**Fig. 8**). Particularly, relatively lower MST values were observed in WLS than in the other two sites, suggesting a stronger deterministic process in WLS in comparison with other sites (**Fig. 8**).

**Fig. 3.**
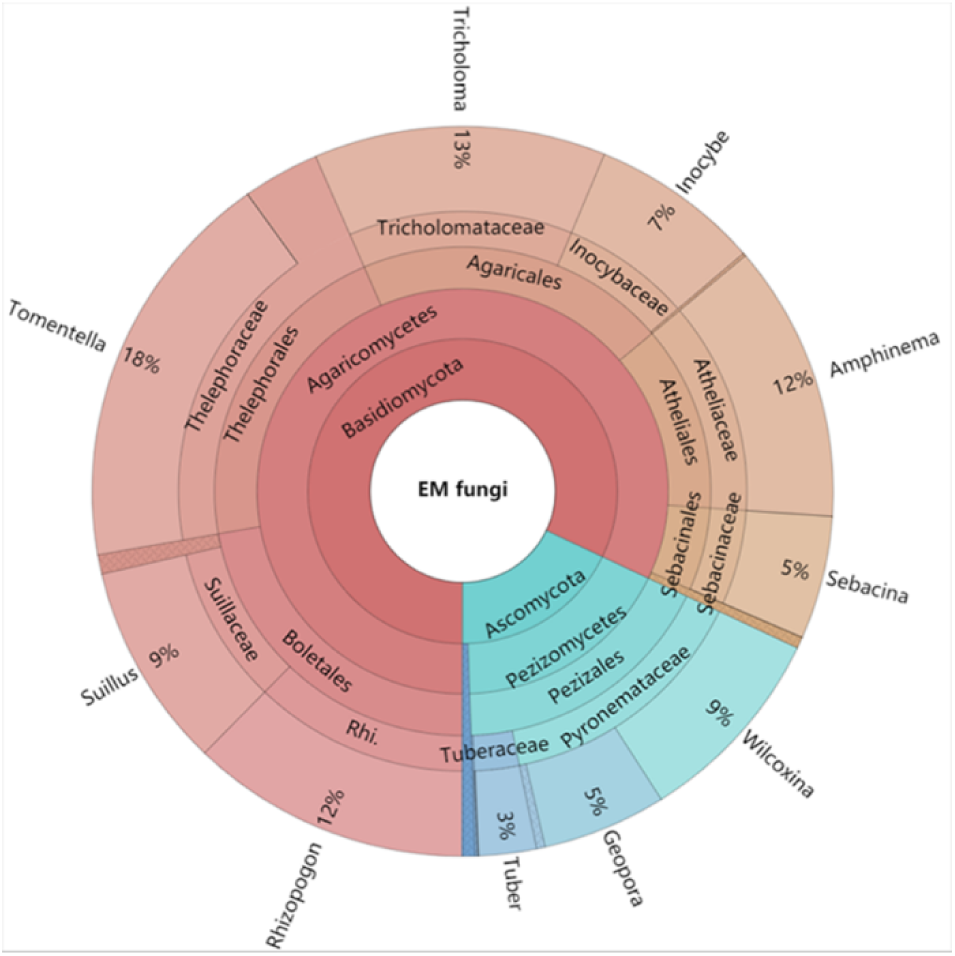
Krona chart of taxonomic affiliation of ectomycorrhizal fungi and their relative abundances. Inner circle represent higher taxonomic ranks and more detailed taxonomic ranks are presented in outer circles. Rhi., Rhizopogonaceae.

**Fig. 4.**
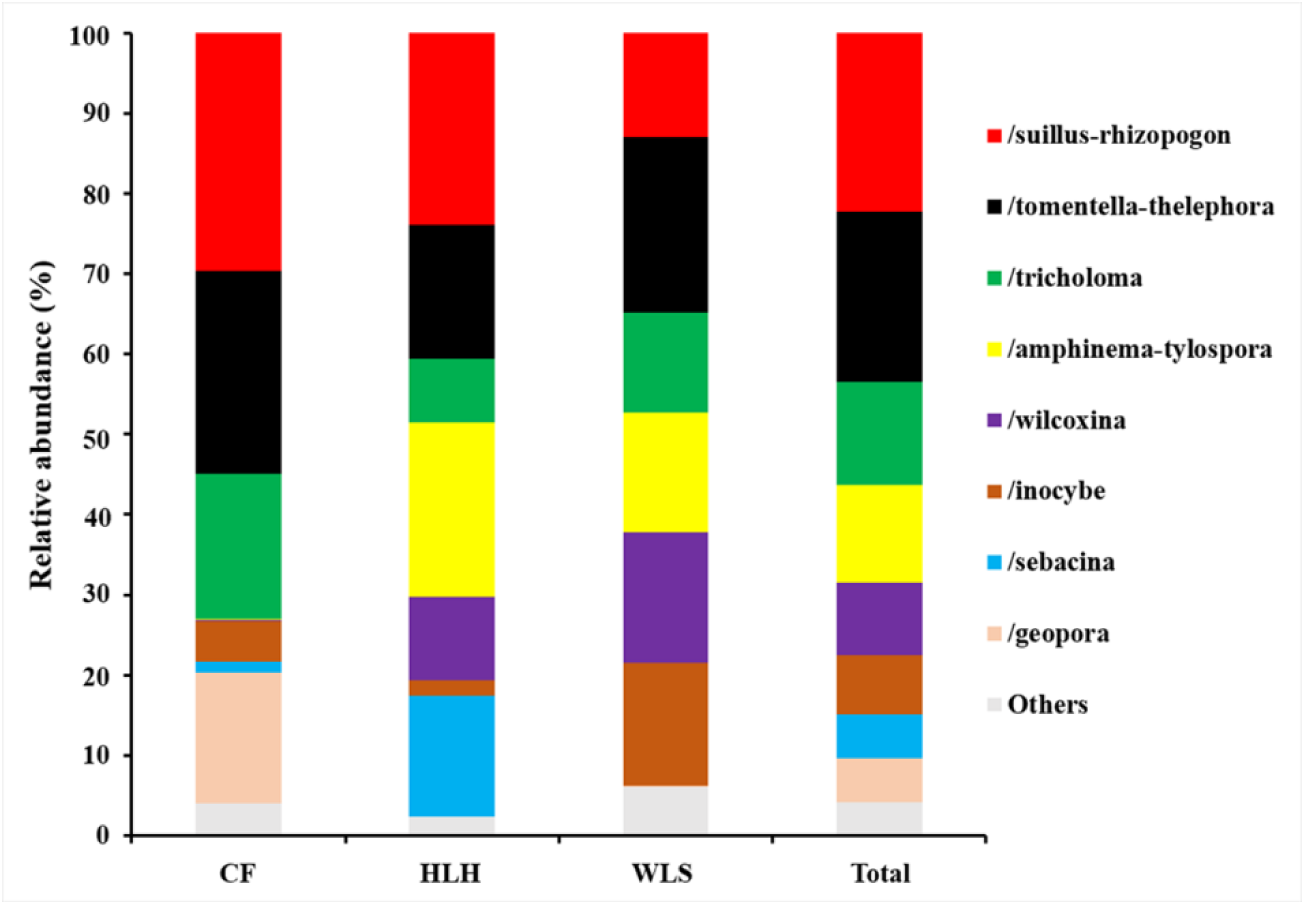
Ectomycorrhizal fungal lineages and their relative abundances. Here only showed the dominant (> 5% of total reads). CF, Chifeng; HLH, Heilihe; WLS, Wulashan; ALL, all samples.

**Fig. 5.**
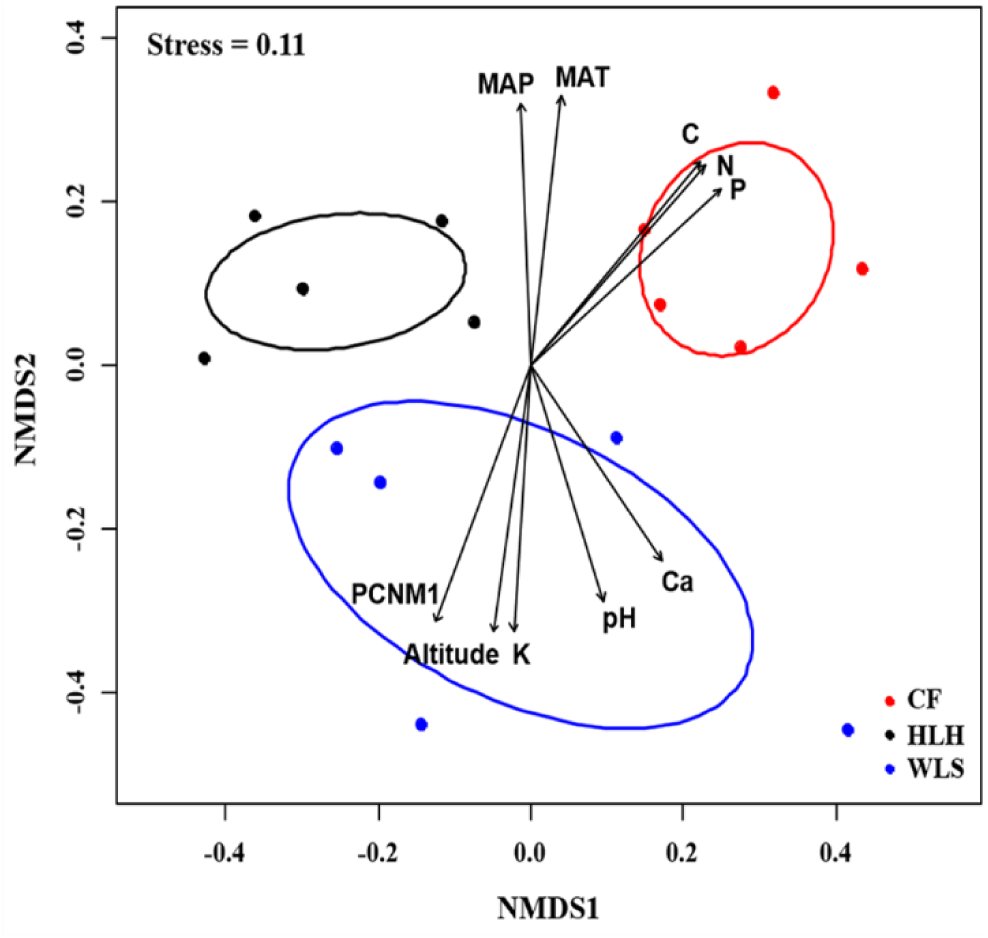
Non-metric multidimensional scaling (NMDS) of the ectomycorrhizal fungal community. Ellipses indicate 95% confidence intervals around centroids for each site. Significant spatial, soil and climatic variables were fitted onto the NMDS ordination. PCNM, principal coordinates of neighbor matrices; MAT, mean annual temperature; MAP, mean annual precipitation; C, soil total carbon; N, soil total nitrogen; P, soil total phosphorus; Ca, soil total calcium; K, soil total potassium.

**Fig. 6.**
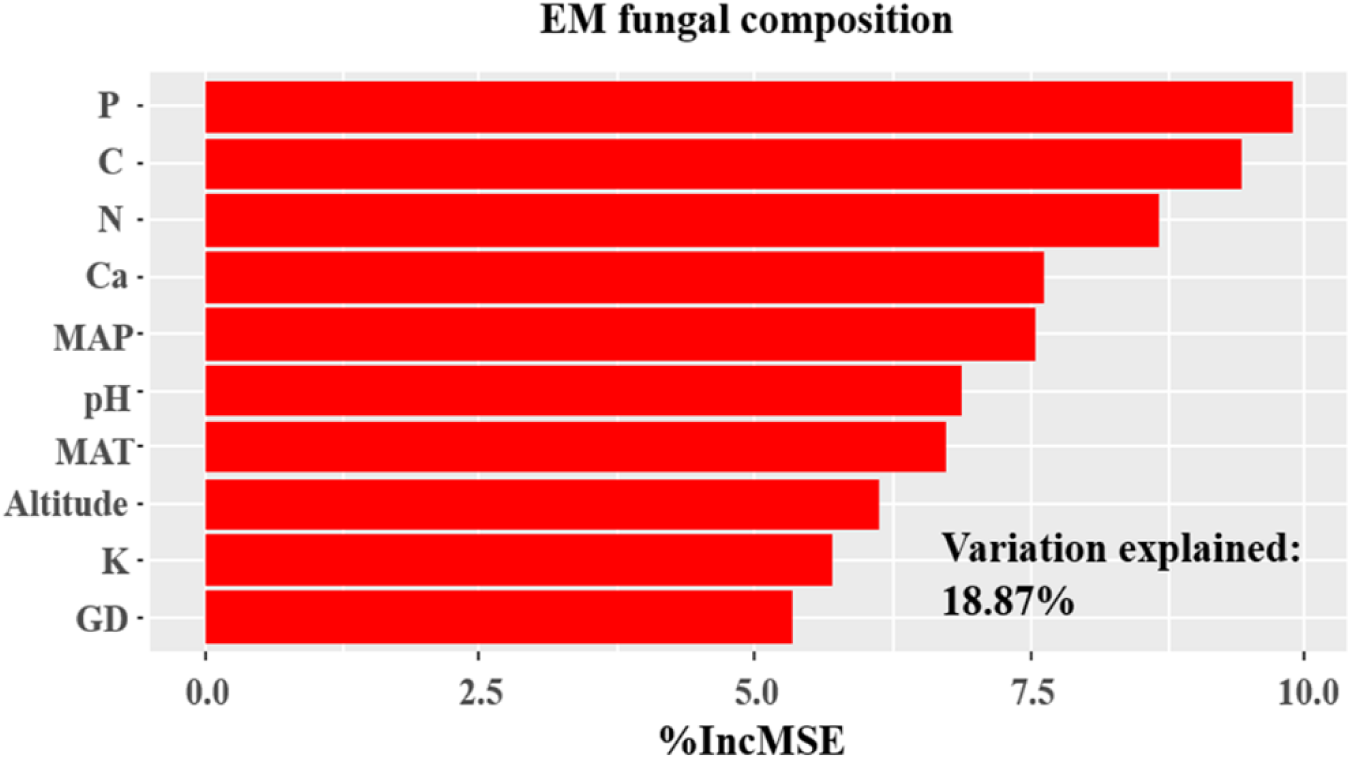
Random forest model showing relative importance of spatial, soil and climatic variables for variations in ectomycorrhizal (EM) fungal community of three sites. %IncMSE, % of increase of mean square error; MAT, mean annual temperature; PCNM, principal coordinates of neighbor matrices; MAT, mean annual temperature; MAP, mean annual precipitation; C, soil total carbon; N, soil total nitrogen; P, soil total phosphorus; Ca, soil total calcium; K, soil total potassium; GD, geographic distance; Significant factors are shown in red.

**Fig. 7.**
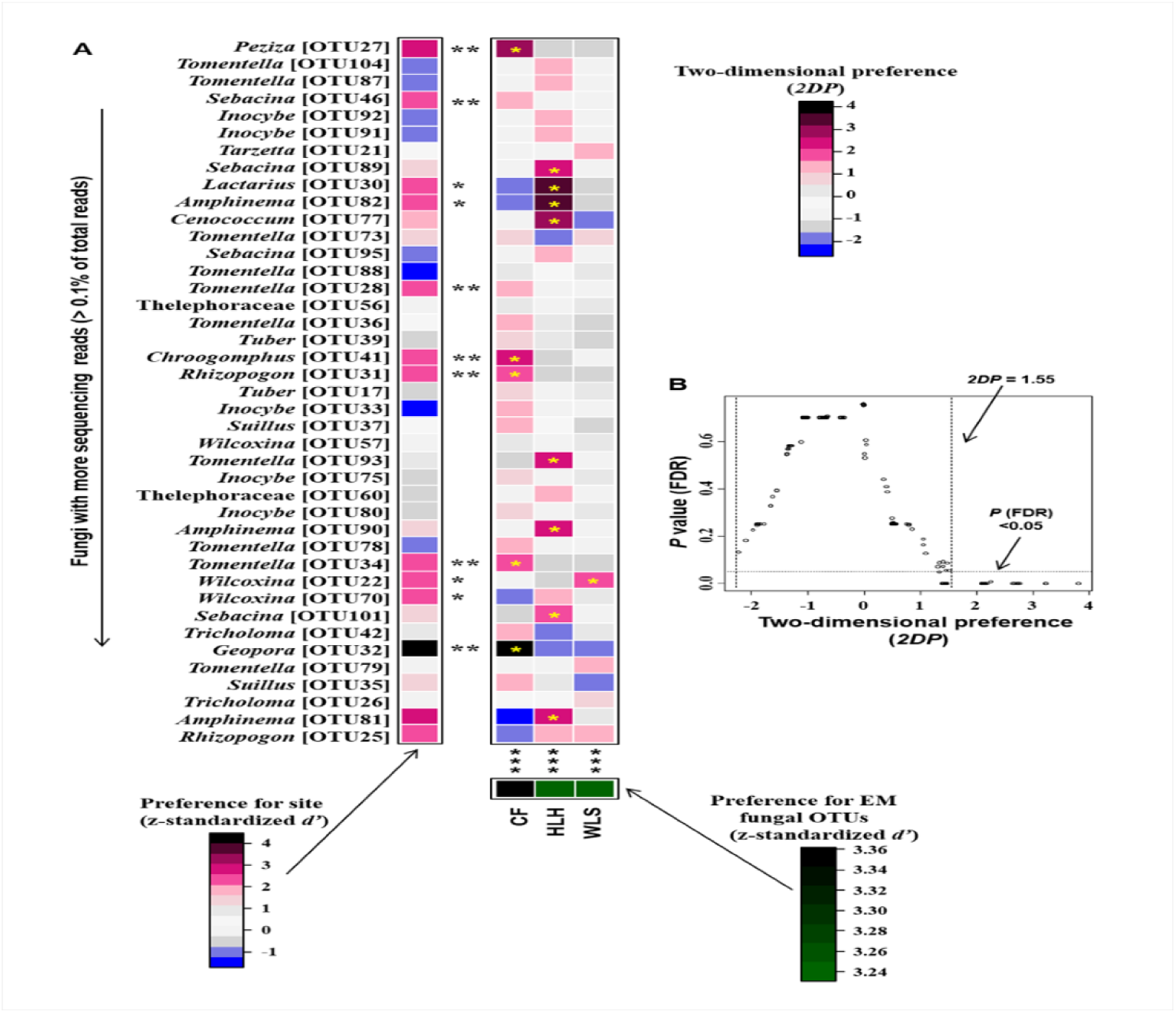
Distribution preferences of site-fungus pairs. (A) standardized *d’* estimates of preferences for fungal operational taxonomic units (OTUs) for indicated sites (columns). Likewise, the standardized *d’* estimate of preferences for sites is indicated for each of the observed fungal OTUs (row). A cell in the matrix indicates a two-dimensional preference (*2DP*) estimate, indicating the extent an association of a focal site-fungus pair was observed more/less frequently than expected by chance. The cell with asterisk inside represents significant preferences in site-fungus pair. Because multiple species/OTUs were tested, the *P* values are shown as false discovery rates (FDRs) in the plant/fungus analysis. (B) relationship between *2DP* and FDR-adjusted *P* values, *2DP* values larger than 1.85 represented strong preferences. Significance: *, *P* < 0.05, **, *P* < 0.01, ***, *P* < 0.001. CF, Chifeng; HLH, Heilihe; WLS, Wulashan.

**Fig. 8.**
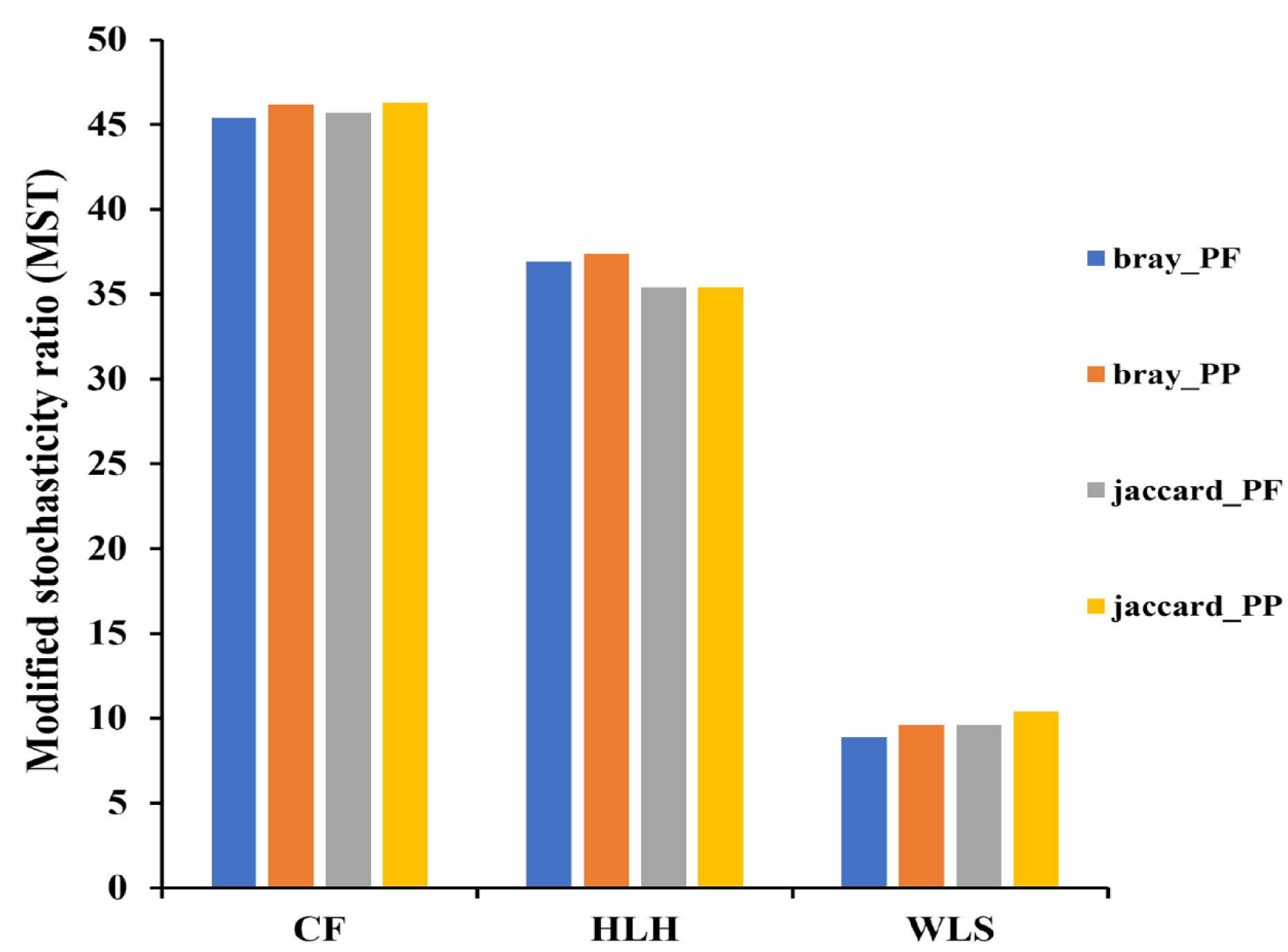
The modified normalized stochasticity ratio (MST) analysis showing the community assembly of ectomycorrhizal fungi. Different colors represent the average MST values calculated based on one distance matrix in couple with one algorithm. CF, Chifeng; HLH, Heilihe; WLS, Wulashan.

**Table 2.**
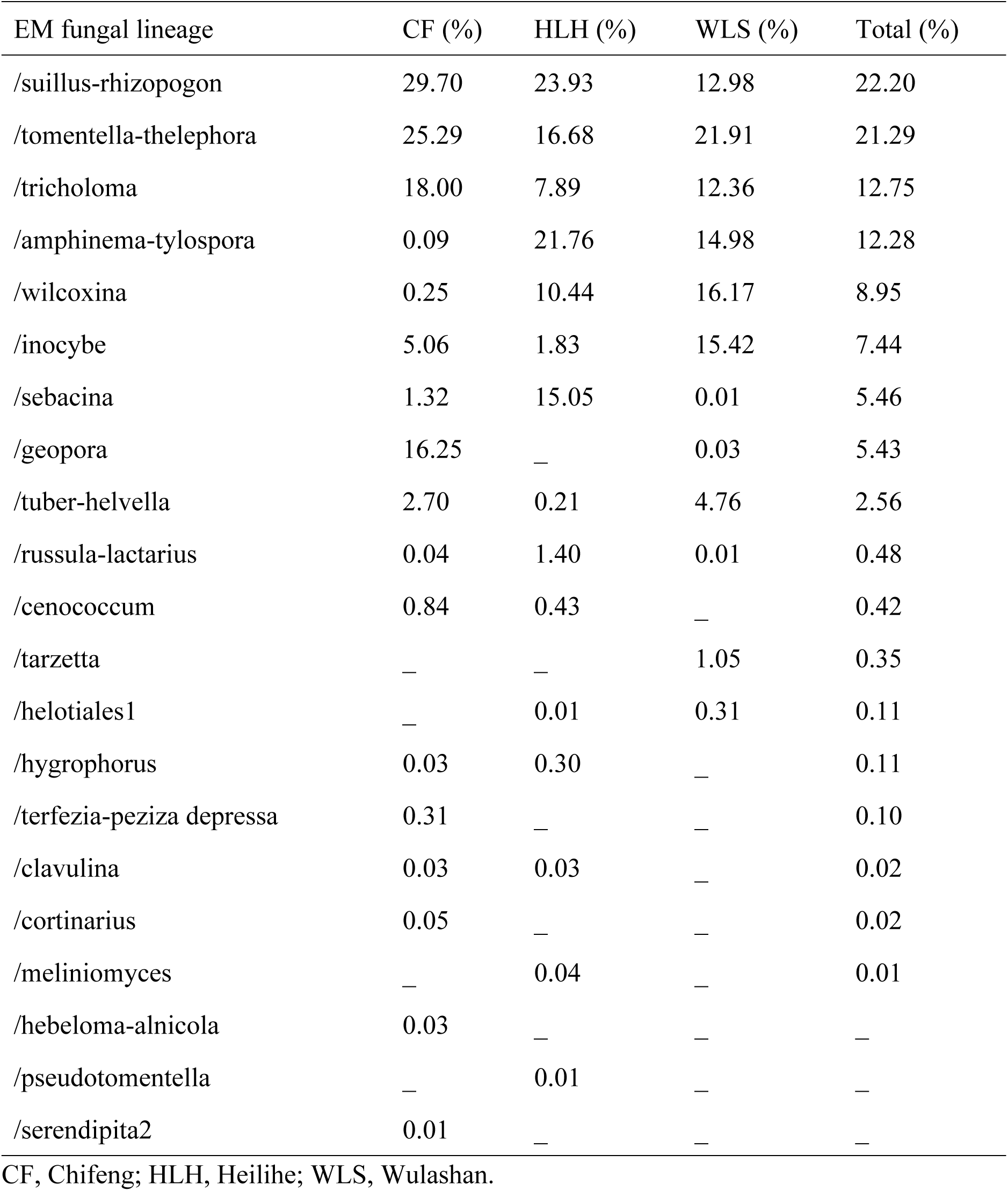
All EM fungal lineages and relative abundances in this study.

| EM fungal lineage | CF (%) | HLH (%) | WLS (%) | Total (%) |
| --- | --- | --- | --- | --- |
| /suillus-rhizopogon | 29.70 | 23.93 | 12.98 | 22.20 |
| /tomentella-thelephora | 25.29 | 16.68 | 21.91 | 21.29 |
| /tricholoma | 18.00 | 7.89 | 12.36 | 12.75 |
| /amphinema-tylospora | 0.09 | 21.76 | 14.98 | 12.28 |
| /wilcoxina | 0.25 | 10.44 | 16.17 | 8.95 |
| /inocybe | 5.06 | 1.83 | 15.42 | 7.44 |
| /sebacina | 1.32 | 15.05 | 0.01 | 5.46 |
| /geopora | 16.25 | — | 0.03 | 5.43 |
| /tuber-helvella | 2.70 | 0.21 | 4.76 | 2.56 |
| /russula-lactarius | 0.04 | 1.40 | 0.01 | 0.48 |
| /cenococcum | 0.84 | 0.43 | — | 0.42 |
| /tarzetta | — | — | 1.05 | 0.35 |
| /helotiales1 | — | 0.01 | 0.31 | 0.11 |
| /hygrophorus | 0.03 | 0.30 | — | 0.11 |
| /terfezia-peiziza depressa | 0.31 | — | — | 0.10 |
| /clavulina | 0.03 | 0.03 | — | 0.02 |
| /cortinarius | 0.05 | — | — | 0.02 |
| /meliniomyces | — | 0.04 | — | 0.01 |
| /hebeloma-alnicola | 0.03 | — | — | — |
| /pseudotomentella | — | 0.01 | — | — |
| /serendipita2 | 0.01 | — | — | — |

CF, Chifeng; HLH, Heilihe; WLS, Wulashan.

## DISCUSSION

A total of 105 EM fungal OTUs were detected in our study, and there was no significant difference of EM fungal OTUs across the three sites, although CF and HLH harbored higher EM fungal diversity than that in WLS. GLM result indicated that MAP, MAT, soil K, and altitude were significant predictors for EM fungal diversity, which has also been reported in previous studies (Wang et al., 2019, 2021). As for the geographic locations of our sampling sites, CF and HLS located in the east of Inner Mongolia, which is the typical temperate forest, while WLS located in the west of Inner Mongolia which is the semi-arid forest. Indeed, the MAT and MAP of WLS were significantly lower than those in CF and HLH. Thus, EM fungal diversity was lower in this arid area as more precipitation could decrease EM fungal diversity (Põlme et al., 2013), accordingly, climatic conditions have been proven to significantly affect EM fungal diversity (Tedersoo et al., 2013a; Miyamoto et al., 2018; Wang et al., 2019b). Additionally, the altitude of WLS was higher than that in CF and WLS, which may be also contributing low EM fungal diversity in WLS due to the fact that the lower energy in higher altitudes could not support high EM fungal diversity, which has been reported in the previous study (Bahram et al., 2011; Gong et al., 2022).

The /*suillus-rhizopogon*, /*tomentella-thelephora*, /*tricholoma*, /a*mphinema-tylospora*, /*wilcoxina*, /*inocybe*, /*sebacina* and /*geopora* were the top eight dominant EM fungal lineages (accounting for 95.8% of total EM fungal sequences), with /*suillus-rhizopogon* and /*tomentella-thelephora* being the most dominant. /*suillus-rhizopogon* has been reported to be specialist for Pinaceae plants. According to Agerer (2001), *suillus* and *rhizopogon* mycorrhizae are typically representatives of the “long-distance exploration type”, which can search for and transport nutrients and water to host plants from a long distance. Due to specificity to certain host plants, *suillus* can connect and transport nutrients to the same plant species, and prevent materials to other tree species (Kennedy et al., 2007). Meanwhile, this lineage was found in most of samples (11 of 15 samples), indicating it is widespread symbiotic partner of *P. tabuliformis* in Inner Mongolia. Our findings suggest /*suillus-rhizopogon* should be an important EM fungal lineage for Pinaceae species. Indeed, the dominance of /*suillus-rhizopogon* has also been reported in studies involved in other Pinaceae species, and has been suspected to help Pinaceae species grow well in harsh habitats (Mandolini et al., 2022; Zhang et al., 2022). /*tomentella-thelephora* has been reported as the dominant EM fungal lineage in many studies conducted on various host plants and diverse forest ecosystems (Tedersoo et al., 2013b), which means that this lineage is widespread group and harbor strong adaptability to environmental changes in the globe. Besides /*suillus-rhizopogon* species host-specialists, there are a number of generalist EM fungi. /*tricholoma* has been reported as the most dominant lineage of *Larix gemelinii* Rupr. in the Great Khingan Mountains, Inner Mogolia (Wang et al., 2021b), which was obviously different in comparison with our study. This may be due to strong host selection on EM fungal communities derived from coevolution between host plant and EM fungi. Correspondingly, strong host effect on EM fungal communities has been intensively investigated in diverse forests and plant identity (Tedersoo et al., 2013a; Wu et al., 2018; Wang et al., 2019; Yang et al., 2022).

NMDS ordination in couple with PerMANOVA indicated that the EM fungal communities differed significantly among the three sites, mirroring a strong spatial effect. This can be explained by the fact that relative abundances of some abundant EM fungal OTUs varied across sites. Meanwhile, Preference analysis indicated that 11 out of 41 (26.8%) abundant EM fungal OTUs showed significant preference to specific site. Additionally, Venn diagrams showed that there were only eight OTUs shared among sites, and 71.4% of total OTUs existed in one site. Thus, the differences in relative abundances, site-preference, and frequency of EM fungi contributed to the significant influence of site on EM fungal communities. In the present study, Random Forest analysis also suggested that geographic distance significantly affected EM fungal community. Thus, these results in combination with NMDS ordination indicated that the fungal community of EM fungi associated with *P. tabuliformis* in Inner Mongolia was spatial structured. This is in accordance with previous studies those implying EM fungi are found to be spatial structed at various scales ranging from local to continental scales (Pickles et al., 2012; Glassman et al., 2015; Wu et al., 2018; Wang et al., 2019b; Wang et al., 2021a, b). This may be due to the dispersal limitation of EM fungi impendent by geographic distance, that is, fungal species failed to disperse from one site to other habitats, or dispersal at a limited distance (Peay et al., 2010; 2012). Environmental variables such as soil and climatic factors were significant predictors for the EM fungal community revealed by environmental fitting test and random forest analysis, which implied a strong effect of environmental filtering on EM fungi; that is, deterministic processes drove community assembly of EM fungi. Accordingly, the results of MST analysis based on two models and two algorithms all supported that deterministic process played more important roles in predicting EM fungal community assembly than stochastic process. We suspect that the dominance of deterministic or stochastic processes on EM fungal community assembly was related with community size (i.e. species richness). Indeed, in our study, EM fungal richness in WLS was lower than that in CF and HLH, and deterministic processes played a stronger role in shaping EM fungal community in WLS than that in CF and HLH. This may be related to interactions among EM fungal species, but this needs further investigation in future study.

## CONCLUSIONS

Our study specially revealed the EM fungal diversity, composition, and mechanisms underlying the community assembly of the *P. tabuliformis* in China for the first time. We found that this Pinaceae plant harbored a high diversity of EM fungi. The /*suillus-rhizopogon* and /*tomentella-thelephora* were the most dominant EM fungal lineages. Most of EM fungal OTUs were unique to one site, and some abundant OTUs preferred to exist in certain cite. Deterministic processes played more important roles than stochastic processes in driving EM fungal community assembly. Our study provides fundamental insights into EM fungal diversity and community structure with a single Pinaceae plant species in Inner Mongolia, China.

## ACKNOWLEDGEMENTS

We thank Wen-Qiang Liu from the Wulashan (WLS) Mountain National Forest Park, Inner Mongolia, and Qi Wang from the National Chinese pine(*Pinus tabulaeformis*) seed base of Heilihe (HLH) Forest Farm for help during fieldwork.

---

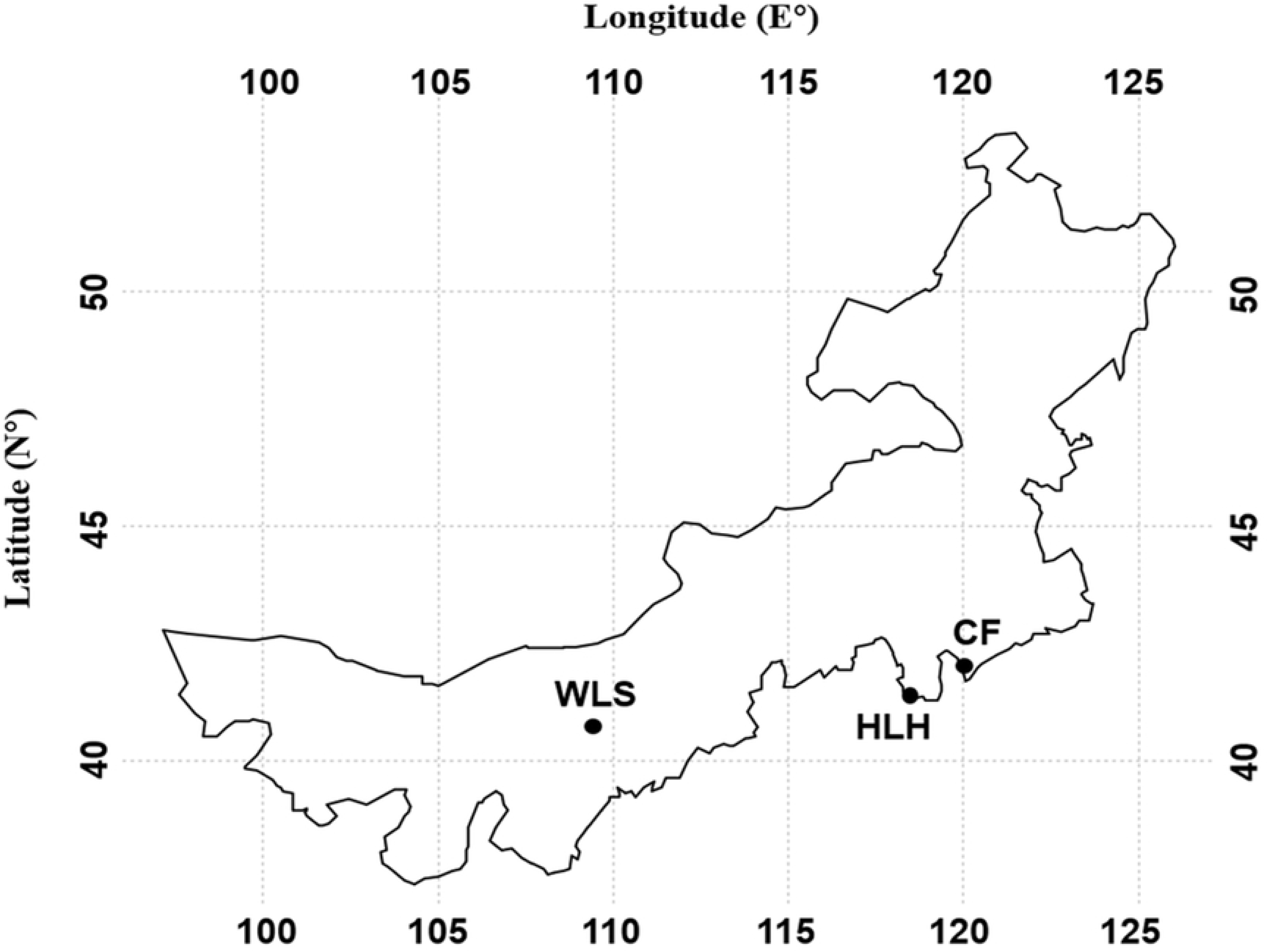

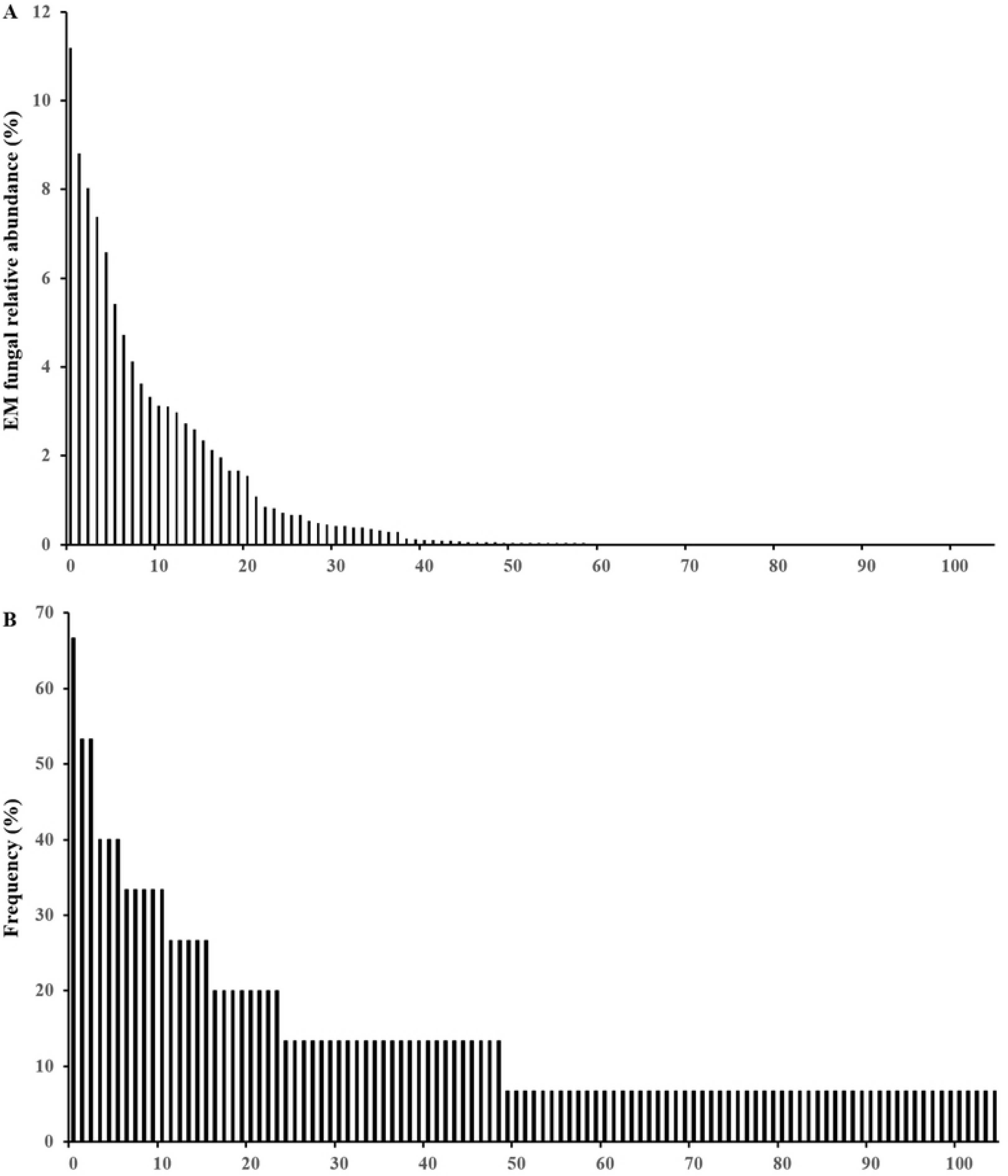

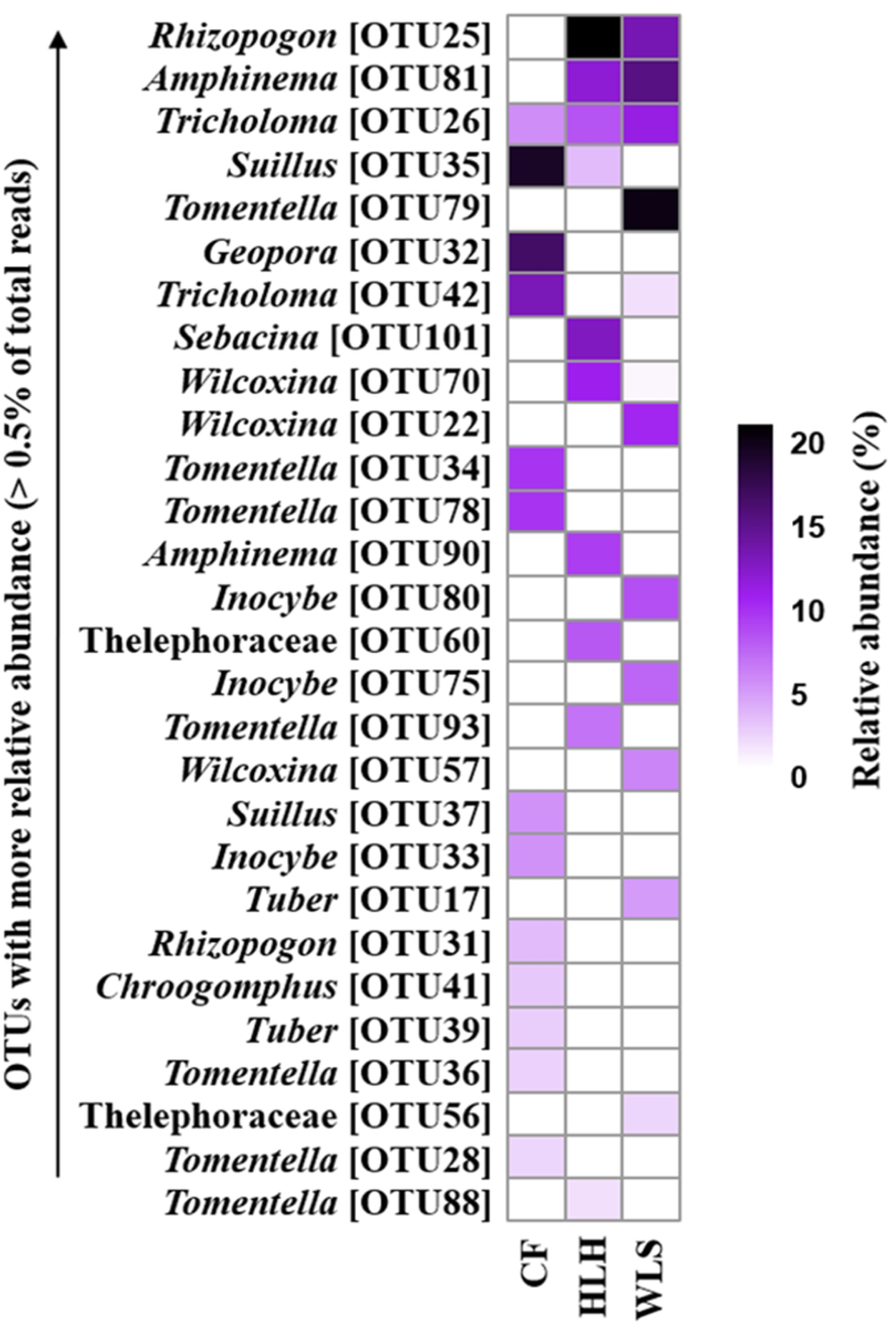

